# Profound taxonomic and functional gut microbiota alterations associated with trichuriasis: cross-country and country-specific patterns

**DOI:** 10.1101/2025.03.21.641387

**Authors:** Pierre H.H. Schneeberger, Julian Dommann, Nurudeen Rahman, Eveline Hürlimann, Somphou Sayasone, Said Ali, Jean Tenena Coulibaly, Jennifer Keiser

## Abstract

**Background:** The human gastrointestinal microbiota plays a crucial role in immune modulation, metabolism, and pathogen resistance. Soil-transmitted helminth (STH) infections, including *Trichuris trichiura* (whipworm), significantly alter gut microbial composition, yet the extent and functional consequences of these changes remain underexplored across different geographical regions. This study investigates the taxonomic and functional impacts of *T. trichiura* on gut microbiota in three endemic regions - Côte d’Ivoire, Laos, and Tanzania - using a unified high-resolution metagenomic sequencing approach.

**Results:** This study reveals extensive gut microbiota disruptions linked to *T. trichiura* infection, with both regional and cross-country patterns. A core signature found across all study sites includes depletion of *Faecalibacterium prausnitzii* and *Eubacterium rectale* (Short-chain fatty acids (SCFA) producers) and enrichment of mucin-degrading bacteria (*Ruminococcus*, *Bacteroides*), alongside increased host-derived carbohydrate metabolism. Infection destabilized microbial networks, characterized by reduced connectivity and clustering, with opportunistic taxa such as *Segatella copri* emerging as network hubs across regions, indicating a shared ecological response. Both taxonomic and functional disruptions exhibited a combination of conserved and region-specific patterns; for example, whereas certain taxa, such as *Prevotella* and *Streptococcus*, showed notable geographic variability, specific functional changes, such as SCFA depletion and mucin degradation, were consistently observed across sites. These conserved functional changes suggest that *T. trichiura* imposes similar metabolic pressures on the gut microbiome across populations, potentially affecting host nutrient availability and immune responses in a predictable manner.

**Discussion and Conclusion:** This study provides robust evidence that *T. trichiura* infection induces significant and consistent microbiome alterations across diverse populations. The depletion of SCFA-producing bacteria and the enrichment of mucin-degrading taxa, along with corresponding metabolic pathways, imply compromised gut barrier integrity, providing insights into the complex inflammatory processes associated with this helminth infection. Additionally, these microbiome differences could play a critical role in facilitating parasite persistence and reinfection, which remain major challenges limiting the efficacy of global control strategies. Our findings highlight the potential of microbiome-targeted interventions, such as probiotic supplementation or dietary modifications, to mitigate the health impacts of *T. trichiura* infections by restoring microbial homeostasis.

## Introduction

The human gastrointestinal tract harbors a complex and diverse microbial community integral to maintaining health and homeostasis [1]. These microbes engage in essential processes such as nutrient metabolism, immune modulation, and protection against pathogenic invasion. Disruptions to this delicate ecosystem, known as dysbiosis, have been linked to a wide array of health conditions, including metabolic syndromes, inflammatory diseases, and infections [2]. Parasitic infections, particularly those caused by soil-transmitted helminths (STHs), are significant drivers of dysbiosis, profoundly altering gut microbial structure and function [3-6]. Among these helminths, *Trichuris trichiura* - commonly referred to as the human whipworm - remains a critical public health concern due to particularly low treatment success compared to other helminth species, especially in low- and middle-income countries [7, 8]. Globally, it is estimated that more than 477 million people are infected with *T. trichiura*, with the heaviest burden in tropical and subtropical regions where poor sanitation facilitates transmission [8]. The parasite primarily colonizes the large intestine, where it anchors in the colonic epithelium and mucosa, triggering localized inflammation and immune responses that can alter gut microbiota composition [9]. Research on *T. trichiura* infection has highlighted its capacity to influence the microbial community in multiple ways, while the gut microbiota, in turn, plays a role in shaping the parasite’s life cycle. First, the parasite induces a pronounced Th2-mediated immune response characterized by increased mucus production and altered epithelial barrier function, which can create a niche favourable to certain microbial taxa [10]. Second, the parasite’s nutrient competition and physical presence in the intestinal lumen may disrupt microbial interactions, contributing to compositional and functional shifts in the microbiota [11]. Third, *T. trichiura* infection often leads to broader physiological changes, such as altered bile acid profiles and reduced short-chain fatty acid (SCFA) production, further shaping microbial ecology [12]. Conversely, the gut microbiota also plays a crucial role in *T. trichiura* colonization, as microbial communities contribute to egg hatching and larval development, establishing a bidirectional relationship between the parasite and the intestinal microbiome [13]. This bidirectional interplay underscores the complex relationship between the helminth and the gut microbiota. Previous studies have begun to explore the relationship between *T. trichiura* infection and gut microbiota, reporting both increased and decreased microbial diversity depending on the population studied and the specific metrics used [14]. For example, a study using a mouse model of *T. muris* infection suggest that the presence of the parasite can enrich for specific taxa, including *Lactobacillus* species [15], while another study reports declines in beneficial commensals associated with *T. trichiura* infection in a small human cohort [15]. Despite these findings, substantial knowledge gaps remain. There is limited understanding of how *T. trichiura* affects microbial diversity and function across diverse geographic regions with varying environmental, dietary, and host genetic factors. Functional consequences of these microbiota changes are also underexplored. While a few studies have highlighted alterations in metabolic pathways, such as reduced Short-chain fatty acids (SCFA) production and shifts in carbohydrate and amino acid metabolism, these findings are often based on single populations or limited datasets. Furthermore, the extent to which *T. trichiura*-induced microbial changes are conserved or vary across regions remains unclear, especially given the significant role that local factors play in shaping the gut microbiota.

This study aims to address these gaps by systematically characterizing the taxonomic, functional, and ecological impacts of *T. trichiura* infection on the gut microbiota across multiple countries using a unified technical framework. By focusing on *T. trichiura* - the only large intestine-colonizing helminth species - this work seeks to provide a more comprehensive understanding of the interplay between parasitic infections and microbial ecology. These insights are critical for advancing our knowledge of host-microbiota-parasite interactions and may inform the development of targeted therapies to mitigate the impact of helminth infections on gut health.

## Methods

### Sample Collection and Microscopy

Stool samples were collected as part of a multi-country randomized controlled trial evaluating the efficacy and safety of albendazole-ivermectin combination therapy against *T. trichiura* and concomitant helminth infections [16]. The trial is registered on ClinicalTrials.gov (NCT03527732, registered on 17 May 2018). Stool samples were analyzed using the standard Kato-Katz microscopy method, previously described in the literature [17]. For molecular analysis, a small aliquot (<1 g) of stool was transferred to a 2 mL screwcap cryotube using a UV-sterilized plastic spatula and immediately frozen at −20°C. At the conclusion of the respective trial stage, frozen samples were shipped to Swiss TPH (Allschwil, Switzerland) on dry ice and stored at −20°C until further analysis.

### DNA Isolation

DNA was extracted from stool samples using the QIAGEN DNEasy PowerSoil Pro (Qiagen, Hilden, Germany) following the manufacturer’s protocol. DNA concentration was quantified using 2 µL of extracted DNA with the Qubit 4.0 fluorometer and high-sensitivity DNA quantification kits (Thermo Fisher Scientific, Waltham, MA, USA).

### Shotgun Sequencing

Shallow shotgun sequencing was performed as described previously [18], generating 2.8E+06 to 5.3E+07 reads per sample. Libraries were prepared using the NEBNext Ultra II FS DNA Library Prep Kit (New England Biolabs, Ipswich, MA, USA) according to the manufacturer’s instructions. Final libraries were quantified and pooled equimolarly before loading onto an Illumina NovaSeq 6000 platform. Sequencing was conducted on S4 flowcell (300 cycles) in paired-end mode (2 × 150 bp). Sample-specific sequencing depth is detailed in **Supplementary Table 1**.

### Taxonomic and Functional Profiling

Taxonomic profiling of shallow shotgun sequencing data was conducted with MetaPhlAn v4.1.1 (11 Mar 2024) [19]. The CHOCOPhlAn species-marker database vOct22_CHOCOPhlAnSGB_202403 (October 2022) was used, and mapping was performed with Bowtie2 v2.5.4 [20]. The resulting taxonomic relative abundance tables were merged using the built-in “merge_metaphlan_tables.py” and were filtered for low counts (minimum count = 100; prevalence = 20% of samples) and low variance (50% features removed based on interquartile range) using MicrobiomeAnalyst v2.0 [21]. Functional profiling was performed using HUMAnN4 v4.0.0.alpha.1 with default settings and the UniRef 2019_06 protein database. The unstratified functional abundance tables, including the “reactions” (= abundance tables based on the Enzyme Comission (EC) numbers) tables as well as the “pathabundance” (= abundance tables summarizing the abundance of MetaCyc pathways [22] based on EC tables) files were normalized to copies per million (CPM) using the humann_renorm_table.py script before downstream analyses.

### Generation of microbiome-specific metrics

All analyses were conducted in R-studio 2024 using R version 4.4+[23]. Alpha and beta diversity metrics were calculated to assess microbial diversity within and between samples. Alpha diversity, reflecting the richness and evenness of species within individual samples, was evaluated using the vegan R package v2.6-6 [24]. Metrics included species richness (number of species), the Shannon diversity index for richness and evenness, and the Berger-Parker index to measure community domination. Beta diversity, representing dissimilarity between samples, was assessed using Bray-Curtis dissimilarity, calculated through the “vegdist” function in the same package. Non-metric multidimensional scaling (NMDS) was performed to visualize compositional differences in a reduced dimensional space, with ordination coordinates calculated using the “metaMDS” function configured with k = 2 and try = 10,000 iterations. To statistically evaluate differences in community-level composition, PERMANOVA (Permutational Multivariate Analysis of Variance) and PERMDISP (Permutation-Based Multivariate Dispersion) analyses were performed using the “adonis2”, “betadisper”, and “permutest” functions. The core microbiome, comprising taxa consistently present across a defined proportion of samples, was identified using the “core” function from the microbiome R package v1.23.1 [25]. Taxonomic profiles were visualized with a heatmap generated through an in-house script combining multiple R packages. The ComplexHeatmap package v2.18.0 [26] was used for heatmap generation, with hierarchical clustering performed using the “hclust” function and Ward’s algorithm, and optimized for clarity using the “dendsort” function. The species network correlation analysis was conducted using functions from the SpiecEasi package v1.1.1 [27], the RCy3 package v3.20 [28], and the igraph package v2.0.3 [29]. Network statistics were calculated using the following functions from the igraph package: “graph.density” for network density, “average.path.length” for the average path length, “diameter” for the network diameter, and “transitivity” for the global clustering coefficient. Centrality measures, including degree, betweenness, closeness, and eigenvector centrality, were calculated using the “degree”, “betweenness”, “closeness”, and “eigen_centrality” functions, respectively.

### Statistical Analysis

Group comparisons were conducted using Mann-Whitney U tests for two-group comparisons and Kruskal-Wallis tests for comparisons involving three or more groups, and all *P*-values were adjusted using the Bonferroni family-wise correction [30]. Species prevalence comparisons were performed in R using the “fisher.test” function, and the resulting *P*-values were adjusted using the Benjamini-Hochberg (BH) procedure [31]. To compare bacterial species abundances, several statistical methods and tools were employed. Adjusted Kruskal-Wallis tests and the LEfSe package [32], available on the MicrobiomeAnalyst platform, were used on country-specific subsets to assess enrichment or depletion of taxa associated with infection status. Additionally, MaAsLin2 v1.18.0 [33] was applied to the complete dataset, controlling for the “Country” variable. All *P*-values resulting from these analyses were adjusted using the BH procedure. To minimize false-positive signals, only species with significant corrected *P*-values across all three statistical methods were considered enriched or depleted. Effect sizes of these differences were visualized using log2 fold-change values generated by MaAsLin2. For comparisons of functional profiles, including ECs and MetaCyc pathway abundances, MaAsLin2 was used to assess statistical differences.

### Visualizations

Most visualizations were created and optimized using OriginPro 2024 v10.1.0.178 (OriginLab Corporation, Northampton, MA, USA). Heatmaps were generated directly in R using the ComplexHeatmap package, while Venn diagrams were created using the ggplot2 package v3.5.1.

## Results

### Sampling results and study characteristics

Between September 2018 and December 2019, individuals from the Pujini and Dodo shehias (health districts) on Pemba Island, Tanzania; the Dabou and Jacqueville districts in the Grand Ponts region, Côte d’Ivoire; and the Nambak district in the Luang Prabang province, Laos, were screened for *T. trichiura* as described elsewhere [16]. A total of 8,866 participants were assessed for eligibility. Of these, 160 participants who tested negative for *T. trichiura* (60 in Côte d’Ivoire, 50 in Laos, and 50 in Tanzania) were selected to represent the “uninfected” microbiome group in this study. 632 *T. trichiura*-positive participants were selected to represent the *Trichuris*-infected group (200 in Côte d’Ivoire, 239 in Laos, and 200 in Tanzania) and were included in this study (**Table 1**, **Figure 1**). 9/799 samples included in this study were dropped due to insufficient sequencing quality. Participants were predominantly young, with mean ages ranging from 14.1 years in Tanzania to 24 years in Laos, and an overall age range of 6–60 years across all regions. Females accounted for roughly half of the infected group, with proportions varying slightly between regions (50.5% in Côte d’Ivoire to 56.6% in Tanzania). *T. trichiura* infections were mostly of light intensity, with geometric mean eggs per gram (EPG) values ranging from 380 in Laos to 497 in Côte d’Ivoire, while moderate and heavy infections were less common.

**Figure 1.**
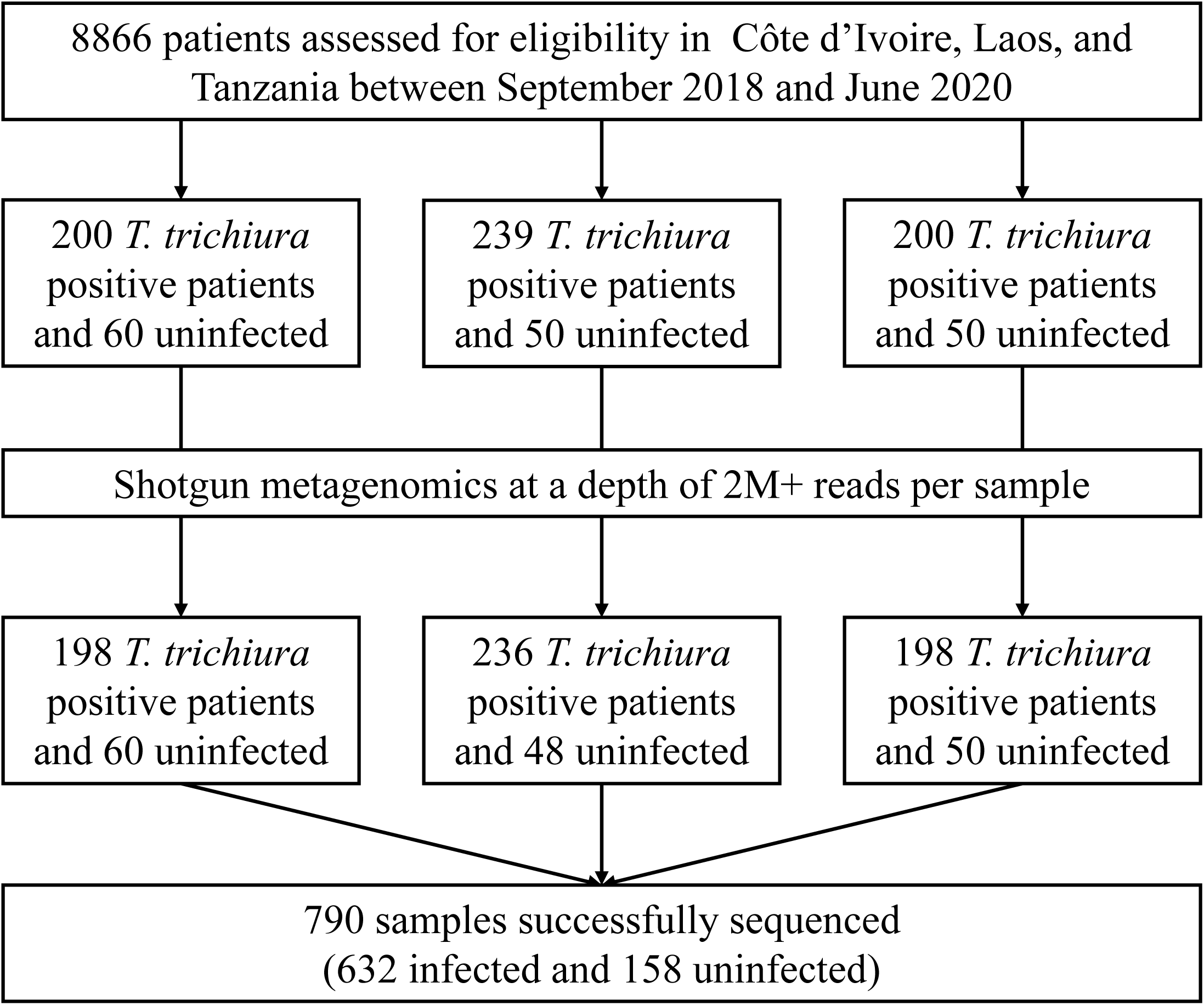
Study profile.

**Table 1.** Cohort description. y = year; EPG = eggs per gram of stool; SD = standard deviation; arit = arithmetic mean; geom = geometric mean; Infection intensity classification is based on World Health Organization criteria [34].

| Characteristic | Côte d’Ivoire | Laos | Tanzania |
| --- | --- | --- | --- |
| **Number of patients (n)** |  |  |  |
| Infected | 198 | 236 | 198 |
| Uninfected | 60 | 48 | 50 |
| Total | 258 | 284 | 248 |
| **Age (y)** |  |  |  |
| Mean (arit) (± SD) | 16.9 (± 14) | 24 (± 16.4) | 14.1 (± 10) |
| Range | 6 to 60 | 6 to 60 | 6 to 60 |
| **Female, n (%)** |  |  |  |
| Infected | 100 (50.5%) | 125 (53%) | 112 (56.6%) |
| ***Trichuris trichiura* infection** |  |  |  |
| EPG<sub>arit</sub> (± SD) | 1081 (± 2047) | 709 (± 1255) | 962 (± 1784) |
| EPG<sub>geom</sub> (± SD) | 497 (± 3) | 380 (± 2.7) | 450 (± 3.1) |
| Range (EPG) | 102-15072 | 102-12822 | 18-12750 |
| Light, n (%) | 151 (76%) | 197 (83%) | 28 (71.8%) |
| Moderate, n (%) | 44 (22%) | 38 (16%) | 45 (23%) |
| Heavy, n (%) | 3 (1.5%) | 1 (0.4%) | 3 (1.5%) |

### Community level taxonomic comparisons

A comparison of alpha diversity metrics between infected and uninfected individuals revealed significant differences across the three countries, as shown in **Figure 2A**. Richness was significantly higher in infected participants from Laos (*P*<0.001) and Tanzania (*P*=0.001), whereas infected study participants in Côte d’Ivoire exhibited significantly lower richness compared to uninfected individuals (*P*<0.001). A similar trend was observed for composite metrics such as Shannon Diversity; however, this difference was not statistically significant for the Tanzanian cohort (*P*=0.11). Analysis of dominance, using the Berger-Parker index, indicated significantly elevated dominance in uninfected individuals in Laos (*P*<0.001), suggesting differing community structures between infection states. Community-level taxonomic composition, evaluated using PERMANOVA (**Figure 2B**), showed that infection status was significantly associated with shifts in community structure in all three countries (*P*<0.001). PERMDISP analysis revealed significant differences in dispersion around the group centroid in Côte d’Ivoire (*P*=0.008) and Laos (*P*=0.049), indicating greater heterogeneity in these settings, while no such differences were detected in Tanzania (*P*=0.472). Core microbiota comparisons (**Figure 2C**) highlighted pronounced differences in microbial composition across countries. Notably, *Bifidobacterium pseudocatenulatum* and *B. adolescentis* were highly prevalent in Tanzania but largely absent in Côte d’Ivoire and Laos, confirming country-specific microbial structures. Clustering of all samples based on species-level abundances (**Figure 2D**) revealed distinct country-specific clusters. While infection status did not lead to discernible clustering in Côte d’Ivoire and Tanzania, a significant association was observed in Laos (Chi-square with Yates correction; *P*<0.001). Specifically, one sample cluster from Laos contained a disproportionately higher proportion of infected patients (n = 90/92), indicating potential infection-related microbial patterns unique to this setting (**Supplementary Figure 1A**).

**Figure 2.**
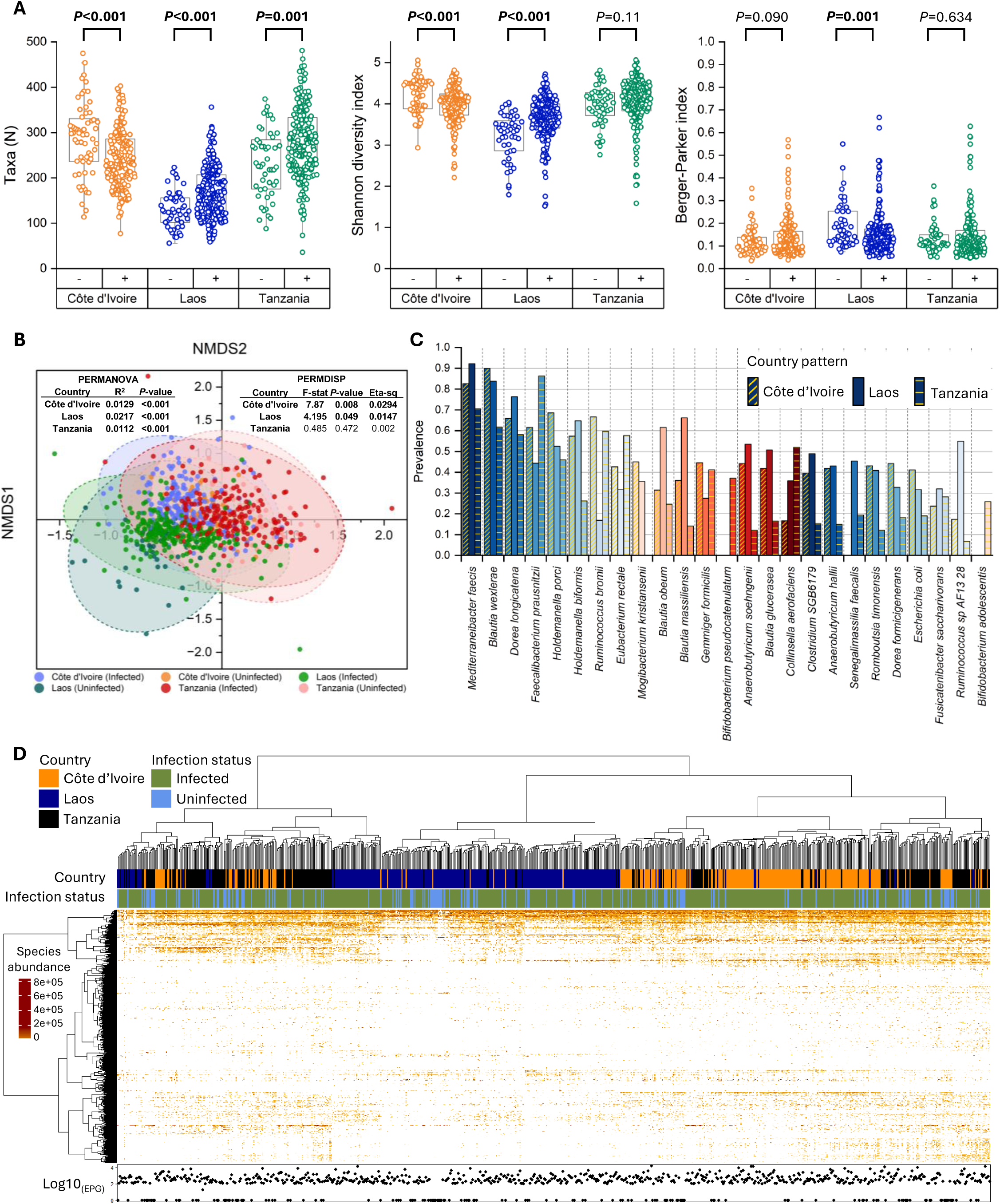
Community-level changes associated with infection status, stratified by country. **Panel A:** Alpha diversity metrics stratified by country and infection status. The left panel shows species richness, the middle panel shows the Shannon diversity index, and the right panel displays the Berger-Parker dominance index. **Panel B:** Non-metric multidimensional scaling (NMDS) plot based on the Bray-Curtis dissimilarity index. Results from community comparisons using PERMANOVA and community dispersion using PERMDISP are shown, with significant results highlighted in bold. **Panel C:** Core microbiome analysis illustrating baseline taxonomic differences across the three study settings. **Panel D:** Heatmap with overall sample clustering based on Robust Aitchison dissimilarity and the Ward method for cladogram construction. The original eggs per gram (EPG) values of stool are displayed at the bottom of each sample.

### Species-level taxonomic signal associated with infection status across countries

A prevalence-based comparison between infected and uninfected individuals, analyzed using a Fisher exact test with Benjamini-Hochberg correction, highlighted significant differences across multiple bacterial taxa (**Figure 3A**). A total of 104 species showed variations in prevalence across the three countries studied. Notably, the genus with the highest number of significant differences was *Streptococcus*, comprising 10 species that were either more or less prevalent depending on infection status. Among these, several species demonstrated consistent differences in prevalence across two or more countries, including *S. sanguinis* (n=2), *S. parasanguinis* (n=2), *S. gallolyticus* (n=2), *S. australis* (n=2), and *S. gordonii* (n=3). In **Figure 3B**, we present modelled species differences based on their relative abundances using a stringent multi-tool statistical approach that combined the Kruskal-Wallis test, LEfSe, and MaAsLin2 (adjusted for country as a variable). Species found to be enriched or depleted in this analysis often overlapped with those identified in the prevalence-based model. For example, several taxa from the *Actinomyces*, *Blautia*, *Clostridium*, *Roseburia*, *Ruminococcus*, *Segatella*, and *Streptococcus* genera showed significant differences in both prevalence and relative abundance models. Interestingly, many species that were differentially abundant exhibited a country-specific fold-change, which was frequently opposite in direction. Further strain-level analysis (**Supplementary Figure 2**) revealed that the divergences were primarily attributable to different strains within the same species, indicating that strain-level variations, rather than species-level differences alone, drive the community shifts associated with *T. trichiura* infection status. In **Figure 3C**, we present the comparison of the cumulative relative abundance of species associated with infection status, stratifying taxa into those enriched or depleted in only one country versus those consistently affected in two or more countries. While there was no significant difference in the cumulative relative abundance of species unique to individual countries, overlapping taxa demonstrated significant differences in cumulative relative abundance across the three countries (P_CI_<0.0001, P_LA_=0.0413, P_TA_=0.0062). These findings highlight the core association of these overlapping taxa with the presence or absence of *T. trichiura*, suggesting a conserved response pattern across distinct geographic regions.

**Figure 3.**
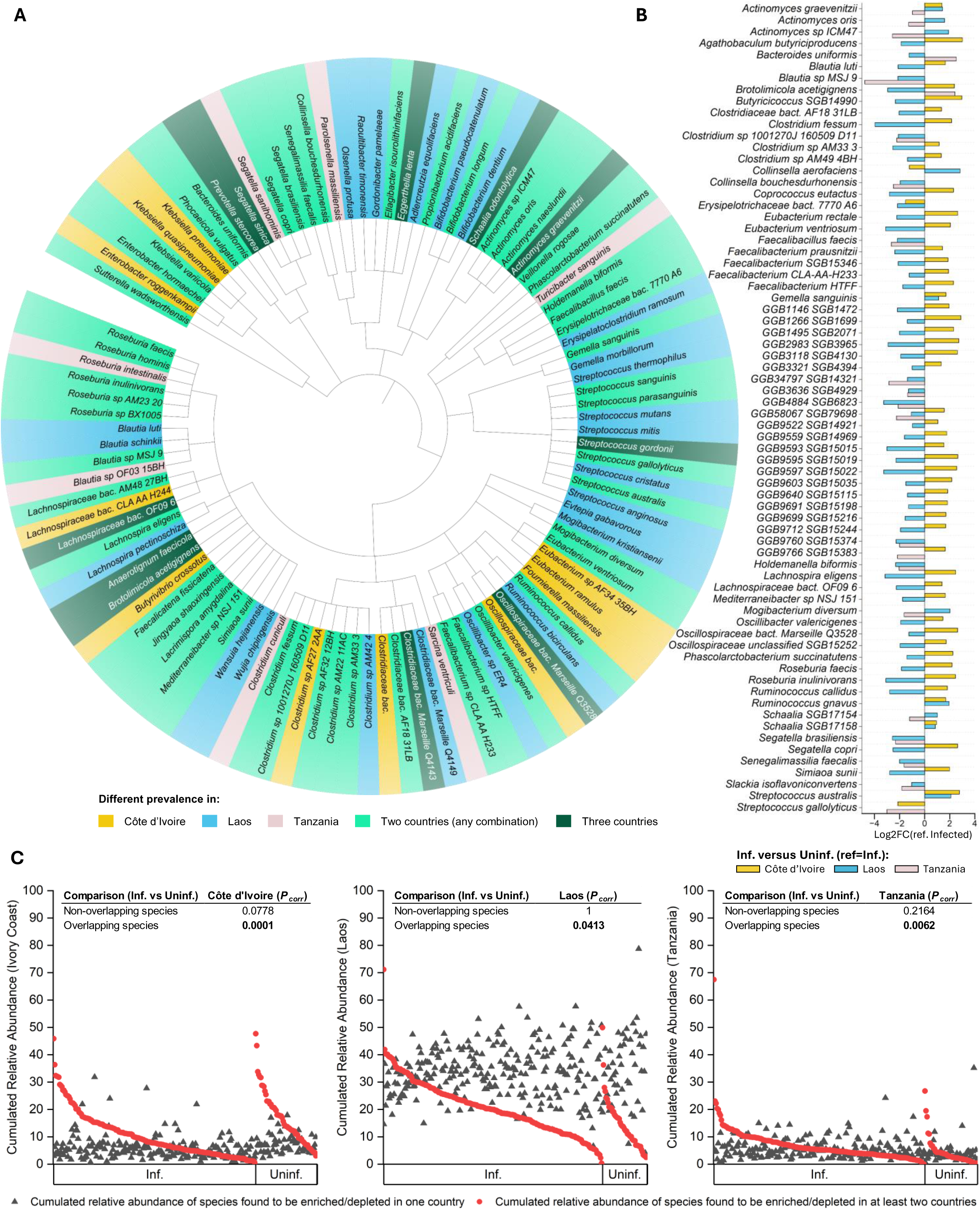
Species-level differences between infection groups, stratified by country. **Panel A:** Prevalence-based comparison showing phylogenetic relationships of species that were significantly more or less prevalent in at least one country. Statistical significance was determined using Fisher’s exact test with Benjamini-Hochberg false discovery rate (FDR) correction (adjusted *P* < 0.05). **Panel B:** Comparison of relative abundances between infection status groups, stratified by country. Three statistical tests were applied: Kruskal-Wallis test, LEfSe differential abundance pipeline, and MaAsLin2 (adjusted for the country variable). Only species that were statistically significant across all three tests were retained. The log2 fold-change values, generated using MaAsLin2, are displayed to represent effect sizes. **Panel C:** Comparison of cumulative relative abundances of species significantly enriched or depleted in one versus two (or more) study sites. Statistical comparisons were performed using the Kruskal-Wallis test, with Bonferroni correction for *P*-values (adjusted *P* < 0.05).

### Changes in species networks associated with infection status

To explore the effects of *T. trichiura* infection on microbial community interactions, we performed network-based analyses using SpiecEasi on taxonomic profiles stratified by infection status and country. Metrics were calculated using the 200 most abundant species, and visualizations focused on the top 50 species in each group (**Figure 4**). Species networks built from infected and uninfected participant samples displayed marked differences in structure across all countries. Key metrics, such as the number of edges (E), network density (De), average path length (PL), diameter (Di), and clustering coefficient (CC), consistently highlighted disrupted microbial interactions associated with infection status. In Côte d’Ivoire, networks from infected participants had an increased number of edges (E=497 vs. 385), higher density (De=0.025 vs. 0.019), and greater clustering coefficient (CC=0.198 vs. 0.147), along with a shorter average path length (PL=3.5 vs. 4.1). These changes suggest a more fragmented yet locally interconnected microbial network. In Laos, infected participants exhibited a lower number of edges (E=306 vs. 368), lower density (De=0.015 vs. 0.018), and shorter average path length (PL=3.9 vs. 4.6), but an increased clustering coefficient (CC=0.186 vs. 0.115). This indicates a more modular and less connected network structure. Finally, in Tanzania, the clustering coefficient was higher in infected participants (CC=0.187 vs. 0.107), while the path length was shorter (PL=4.5 vs. 5.3) and diameter was reduced (Di=13 vs. 17). These changes point to a more compact and tightly clustered microbial network. Infected participant’s networks consistently showed reduced inter-species interactions, with several taxa shifting in their network roles. Genera such as *Streptococcus*, *Clostridium*, *Dorea*, and *Blautia* were central, highly connected nodes in uninfected participant’s networks but became peripheral in infected networks. Conversely, species like *Segatella copri* emerged as a new hub in infected groups, suggesting it’s increased ecological importance or resilience in the context of infection. Some species – including *Faecalibacterium prausnitzii* and *Eubacterium rectale* - became central hubs in infected patients in Côte d’Ivoire and Laos, but not in Tanzania, thus also indicating a widespread, yet locally adapted, ecological response to *T. trichiura* infection. These changes underscore a reorganization of microbial community dynamics in response to *T. trichiura* infection. While infection-induced network perturbations were evident across all countries, their magnitude varied, reflecting the influence of local environmental and host factors. Nevertheless, conserved patterns emerged, including reduced connectivity and clustering in infected networks, supporting the hypothesis that *T. trichiura* infection disrupts microbial community stability.

**Figure 4.**
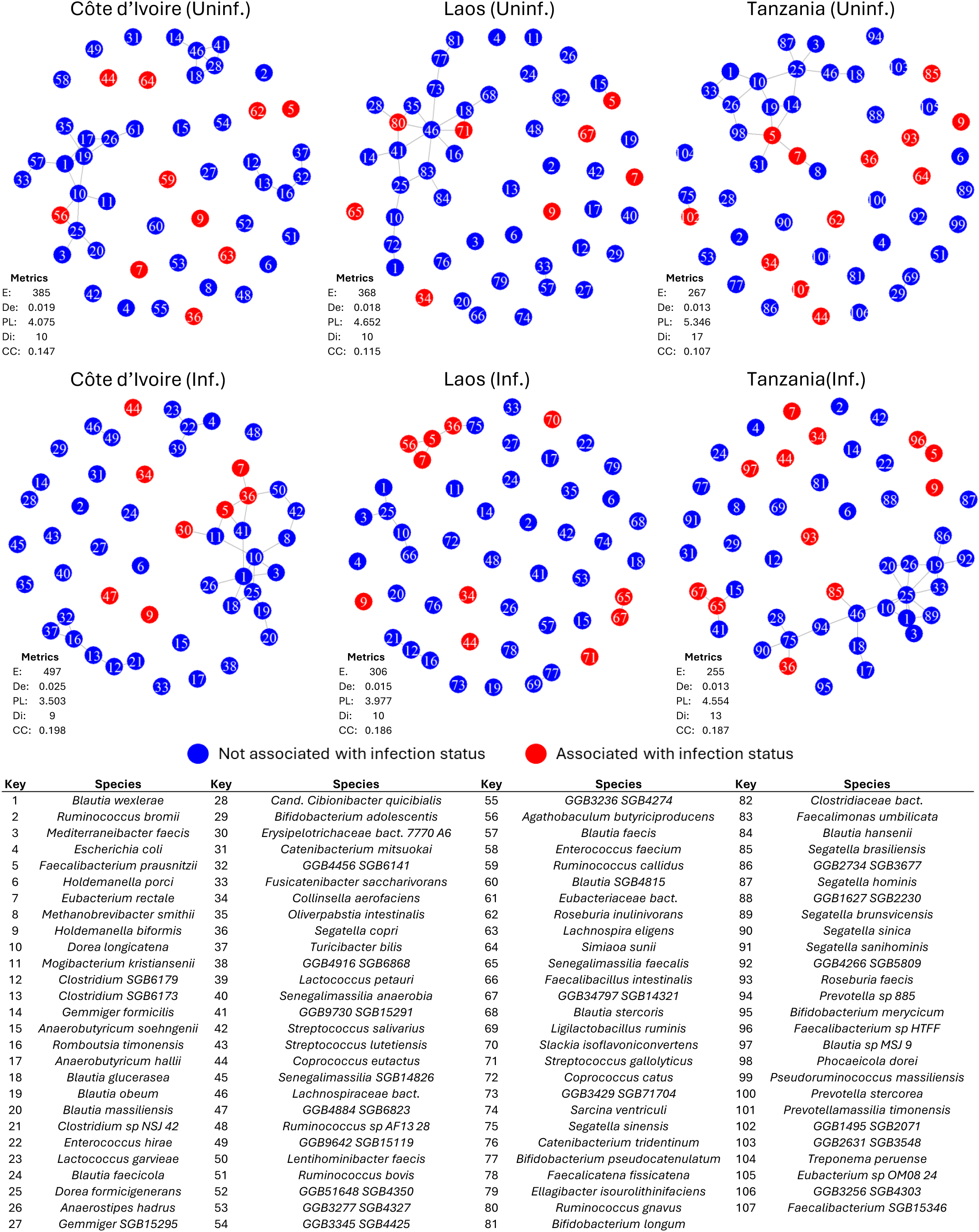
Effect of infection status on the microbiome network correlation structure. SpiecEasi analysis of taxonomic profiles stratified by country and infection status. The visualization on the 50 most abundant species in the different groups, the metrics were calculated using the 200 most abundant species in each group. Network metrics are as follows: E = edges, De = density, PL = average path length, Di = diameter, and CC = clustering coefficient. The numbers in the correlation graphs correspond to the species listed in the table at the bottom of the figure. Inf. = infected; Uninf. = uninfected.

### Functional shifts in microbial communities associated with T. trichiura infection

To investigate functional changes associated with *T. trichiura* infection, we analyzed abundance profiles based on Enzyme Commission (EC) numbers, which systematically classify enzymes by function, across infection status and countries. A total of 6,440 ECs were found, of which 5,453 were shared between the three countries, regardless of infection status (**Supplementary Figure 3A**). As shown in **Supplementary Figure 3B**, functional richness was found to be significantly lower in Laos compared to Côte d’Ivoire (*P*<0.0001) and Tanzania (*P*<0.0001), as well as significantly different when stratified for infection status in Côte d’Ivoire (*P*=0.02) and Tanzania (*P*<0.001) but not in Laos (*P*=0.68). Difference between infected and uninfected groups were observed in each country, with infected groups showing 458, 615, and 840 unique ECs in Côte d’Ivoire, Laos, and Tanzania, respectively (**Figure 5A**). In contrast, uninfected groups exhibited lower unique functions, with 122, 38, and 25 unique ECs in the respective countries, indicating a reduction in function associated with infection status. NMDS analysis based on Bray-Curtis dissimilarity (**Figure 5B**) revealed distinct clustering by infection status and country. PERMANOVA confirmed significant differences between infected and uninfected groups (Côte d’Ivoire: R^2^=0.0076, *P*=0.091; Laos: R^2^=0.0217, *P*=0.001; Tanzania: R^2^=0.0098, *P*=0.014), thus indicating a stronger impact of *Trichuris* infection on the functional potential of gut communities in Laos. PERMDISP analysis indicated no significant differences in community dispersion in Laos and Tanzania (*P*>0.05), but Côte d’Ivoire showed marginal significance (F-stat=3.89, *P*=0.035), suggesting slightly greater variability in functional profiles within infected communities. Volcano plots (**Figure 5C**) highlighted ECs significantly enriched or depleted in infected versus uninfected groups. Overall, the highest number of differentially abundant ECs were found in Laos (n = 1279), followed by Côte d’Ivoire (n = 317) and Tanzania (n = 293). Pathway-level comparisons (**Figure 5D, Supplementary Table 2**) revealed infection-associated shifts in core metabolic processes. We identified significant alterations in 67 MetaCyc pathways between *Trichuris*-infected and uninfected patients using Maaslin2 and controlling for the “Country” variable (*P_adj_*<0.05). Several pathways related to carbohydrate metabolism, fermentation, and biosynthesis were differentially abundant between *Trichuris*-infected and non-infected individuals, reflecting functional reorganization of the gut microbiome. Among the pathways significantly depleted in infected individuals, PWY_5514 (log2_FC_=-0.494, FDR=6.86E-05) and UDPNACETYLGALSYN_PWY (log2_FC_=-0.518, FDR=0.000103), both involved in carbohydrate biosynthesis, exhibited strong reductions, suggesting disruptions in glycan metabolism. PWY_4984, which is involved in inorganic nutrient metabolism, was also underrepresented (log2_FC_=-0.234, FDR=0.000174), indicating altered nutrient utilization. Fermentation-related pathways, many contributing to SCFA metabolism, were also significantly reduced. PWY_3801, linked to mixed-acid fermentation, showed a log2_FC_ of −0.37 (FDR=0.00076), while PWY_7688, a component of carbohydrate biosynthesis, was depleted (log2_FC_=-0.451, FDR=0.0041). In Laos, PWY_7254, a butyrate-producing fermentation pathway, was significantly reduced (log2_FC_=-0.259, FDR=0.0122), suggesting a lower microbial capacity for SCFA production. In contrast, non-infected individuals, particularly in Côte d’Ivoire, exhibited enrichment of fermentation pathways, including PWY_6545 (log2_FC_=0.066, FDR=0.0146) and PWY_6731 (log2_FC_=0.146, FDR=0.0153), which contribute to SCFA biosynthesis. These pathways are essential for gut health and immune modulation, and their depletion in infected individuals may indicate reduced SCFA availability, with potential implications for gut barrier function and host immunity. Finally, several pathways were enriched in non-infected individuals, including PWY_I9 (log2_FC_=0.225, FDR=0.00023), involved in amino acid biosynthesis, and GLUDEG_I_PWY (log2_FC_=0.244, FDR=0.017), associated with amine and polyamine degradation. These differences suggest infection-driven or -associated shifts in nitrogen metabolism.

**Figure 5.**
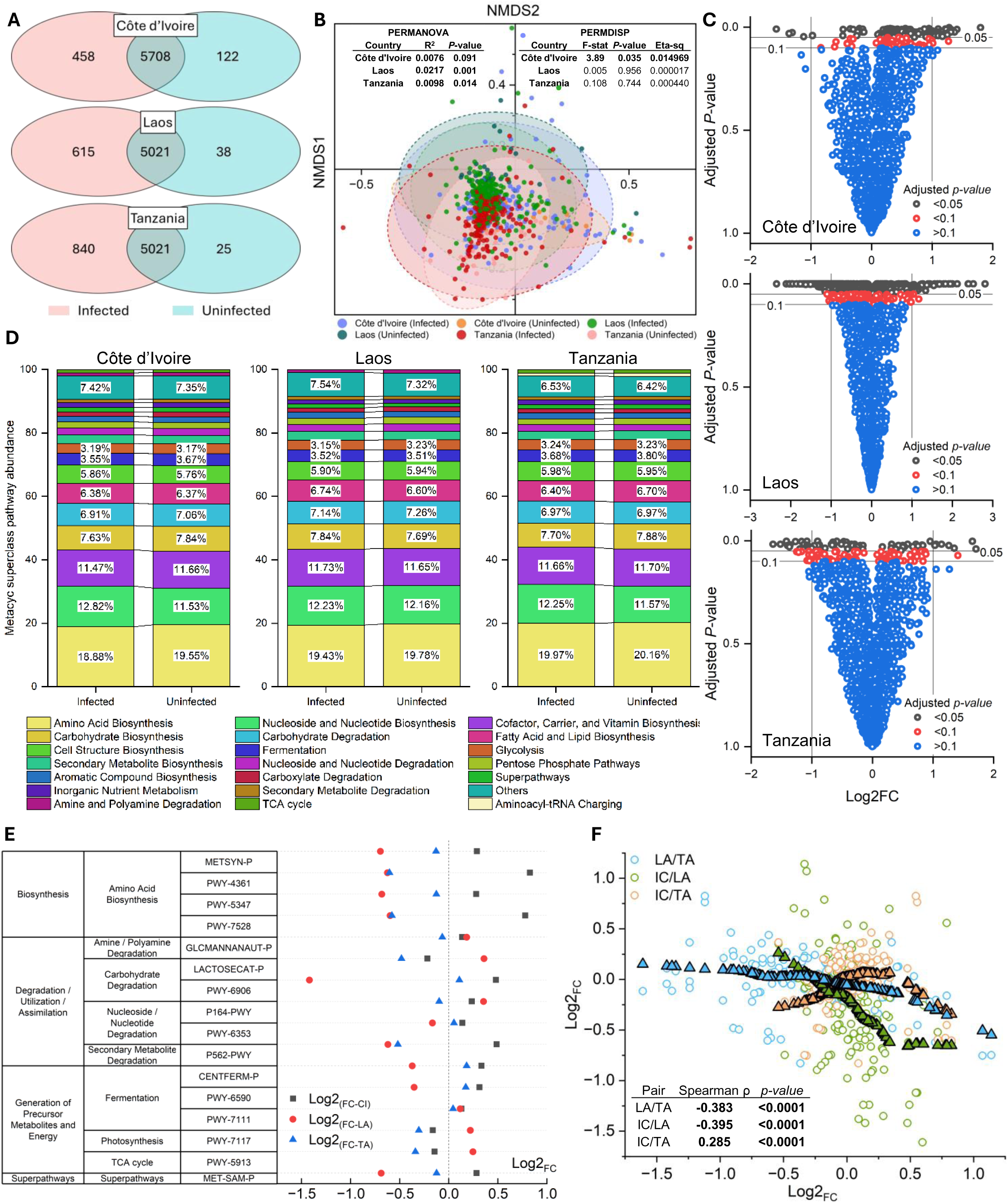
Community-level functional changes associated with infection status, stratified by country. **Panel A:** Venn diagram showing the enzyme commission (EC) numbers found in each study setting, stratified by infection status and showing the overlapping portion. **Panel B:** Non-metric multidimensional scaling (NMDS) plot based on the Bray-Curtis dissimilarity index applied on the EC abundance tables. Results from community comparisons using PERMANOVA and community dispersion using PERMDISP are shown, with significant results highlighted in bold. **Panel C:** Volcano plots based on comparison by infection status of normalized EC abundance tables using MaAsLin2, stratified by country. **Panel D:** Averaged Metacyc superclass abundance comparison by infection status stratified by country. Panel E: Fold-change abundance difference of Metacyc pathways between infected and uninfected patients calculated with MaAsLin2, stratified by country. **Panel F:** Fold-change (FC) comparison of Metacyc pathways associated with infection status between countries. To visualize the monotonic trend, we applied LOWESS smoothing (represented by colored triangles with black outlines) using a proportion of 0.6, balancing smoothness with the capture of local variations. CI = Côte d’Ivoire; LA = Laos; TA = Tanzania.

The stratified analysis across the three study countries identified several pathways exhibiting either consistent or contrasting enrichment/depletion patterns (**Figure 5E**). MET-SAM-PWY, a superpathway involved in methylation processes, was enriched in Côte d’Ivoire (log2FC=0.282, FDR=0.089) but significantly depleted in Laos (log2FC=-0.689, FDR=0.000988), suggesting regional differences in microbial methylation potential. PWY_5913, associated with the TCA cycle, was depleted in infected individuals in Tanzania (log2FC=-0.34, FDR=0.0389) but enriched in Laos (log2FC=0.247, FDR=0.0705), highlighting country-specific adaptations in microbial energy metabolism in response to infection. LACTOSECAT-PWY, involved in carbohydrate degradation, was depleted in Tanzania (log2FC=-0.482, FDR=0.0389) but enriched in Laos (log2FC=0.359, FDR=0.0288), pointing to potential dietary influences on carbohydrate utilization across regions. PWY-6906, which also plays a role in carbohydrate metabolism, was enriched in infected individuals in Côte d’Ivoire (log2FC=0.478, FDR=0.00774) but strongly depleted in Laos (log2FC=-1.42, FDR=1.53E-09), underscoring divergent microbial responses between these two settings. Conversely, some pathways displayed similar patterns of enrichment across countries. P164-PWY, associated with nucleoside and nucleotide degradation, was consistently enriched in Côte d’Ivoire (log2FC=0.234, FDR=0.0203) and Laos (log2FC=0.354, FDR=0.0445), indicating a conserved microbial response to infection. Similarly, PWY-7111, related to fermentation processes, was enriched in infected participants in both Côte d’Ivoire (log2FC=0.128, FDR=0.0708) and Laos (log2FC=0.117, FDR=0.0402). Finally, GLCMANNANAUT-PWY, involved in amine and polyamine degradation, was also enriched consistently in both Côte d’Ivoire (log2FC=0.136, FDR=0.0949) and Laos (log2FC=0.184, FDR=0.00693). Across pathways associated with *Trichuris* infection status in at least one country, the log2_FC_ values were positively correlated between Côte d’Ivoire and Tanzania (Spearman correlation, ρ=0.285, *P*<0.0001), suggesting a degree of consistency in microbiome functional shifts associated with infection in these two regions (**Figure 5F**). However, these values were strongly negatively correlated between Laos and both Côte d’Ivoire (ρ=-0.395, *P*<0.0001) and Tanzania (ρ=-0.383, P<0.0001), indicating a contrasting pattern of pathway enrichment and depletion between these regions.

## Discussion

This study provides the most comprehensive metagenomic investigation to date of *T. trichiura*- associated microbiome alterations, leveraging shotgun metagenomics across three geographically distinct regions. By comparing infected and uninfected individuals, we identified both conserved and region-specific microbiome disruptions, revealing a consistent ecological impact of *T. trichiura* on gut microbial communities. This study extends previous research on helminth-microbiota interactions [14], by demonstrating that *T. trichiura* induces significant taxonomic and functional shifts that transcend geographic variation, providing a broader, comparative perspective on how soil-transmitted helminths influence gut microbial ecology.

One of the most striking findings is the consistent depletion of SCFA-producing bacteria across all study populations. Infected individuals exhibited a marked reduction in butyrate- and acetate-producing taxa, including *Blautia* sp. MSJ 9 and *Holdemanella biformis*, alongside a depletion of key fermentation pathways, such as PWY-7254 and PWY-3801. These SCFAs play a critical role in maintaining colonic health, reinforcing epithelial integrity, and modulating immune responses [35]. Their depletion in infected individuals suggests a potential weakening of gut barrier function, which could have implications for host susceptibility to inflammation and reinfection. Importantly, this study provides the first multi-regional evidence that *T. trichiura* consistently alters SCFA metabolism, regardless of environmental or dietary differences, reinforcing the predictable ecological pressures imposed by this parasite.

This study also demonstrates for the first time that *T. trichiura*’s impact on specific microbial taxa is highly context dependent. While core microbial functions were similarly affected across regions, taxa such as *Prevotella* and *Streptococcus* exhibited region-specific variations in relative abundance, likely reflecting local environmental pressures contributing to shaping microbiome resilience to helminth-induced dysbiosis. Notably, fermentation pathways were strongly depleted in infected individuals from Laos, while in Côte d’Ivoire and Tanzania, some of these pathways remained stable or were even enriched. These findings suggest that while *T. trichiura* exerts a consistent functional impact on the microbiome, the host-microbiota response is also influenced by region-specific dietary and environmental factors, a novel insight that may have implications for future intervention strategies.

The niche-specificity of *T. trichiura* in the large intestine provides further context for these findings. Unlike *Ascaris lumbricoides* and hookworms, which colonize the small intestine, T. trichiura inhabits the cecum and proximal colon [36], microbial environments characterized by high fermentative activity and mucin metabolism [35]. This distinction is crucial, as helminth infections elicit both local and distal immunomodulatory responses, but the effects are notably stronger at the site of infection [37]. This means that stool samples reliably capture microbiome alterations at the infection site [38], thus strengthening the clinical relevance of this study. By focusing exclusively on *T. trichiura*, this study provides the most detailed characterization to date of how a large-intestinal helminth uniquely reshapes gut microbial ecology.

Functional analysis also revealed significant enrichment of mucin degradation pathways in infected individuals, indicative of a microbial shift toward host-derived glycan metabolism. Specifically, consistent enrichment of the GLCMANNANAUT-PWY pathway (autodegradation of glucosamine- and mannose-containing polysaccharides, key mucin components), together with an increased abundance of known mucin-degrading taxa such as *Ruminococcus* and *Bacteroides*, suggests altered mucus dynamics potentially facilitating parasite persistence. These findings align with previous research linking helminth infections to increased mucin degradation [15, 39], but this study is the first to demonstrate these shifts across multiple geographic regions, reinforcing their potential role in *T. trichiura*’s ability to establish chronic infections.

Beyond functional disruptions, network-based microbial interactions provide novel insights into community-level stability in the infected gut. Network analyses revealed reduced microbial connectivity and clustering in infected individuals, suggesting that *T. trichiura* infection leads to a less cohesive and more fragmented microbiome structure. The emergence of opportunistic taxa such as *Segatella copri* as network hubs in infected individuals suggests that these species may exploit helminth-induced disruptions, potentially reinforcing dysbiosis. Notably, similar microbial network disruptions were reported in helminth-infected indigenous populations in Malaysia [14], but this study extends these findings by demonstrating that *T. trichiura*-induced network instability is a globally recurrent feature, independent of geography.

The potential clinical implications of these findings are significant, particularly in regions where *T. trichiura* remains endemic. The depletion of SCFA-producing bacteria and the enrichment of mucin-degrading taxa suggest that *T. trichiura* infection compromises gut barrier integrity and immune homeostasis, potentially increasing the risk of inflammation, enteric co-infections, and reinfection cycles through a microbiome-mediated process. Furthermore, the observed functional shifts may favour parasite persistence, exacerbating chronic infection burdens in endemic settings. These findings highlight the dual role of the microbiota during *T. trichiura* infection: on one hand acting as a protective barrier, and on the other hand potentially facilitating parasite colonization. A better understanding of these dynamics may pave the way for microbiome-based interventions, such as prebiotic or probiotic therapies aimed at restoring microbial stability and enhancing resistance to reinfection.

Nonetheless, this study has some limitations. The cross-sectional design prevents causal inferences about the relationship between *T. trichiura* infection and microbiome changes. Longitudinal studies that track microbiome dynamics during infection, treatment, and reinfection are necessary to establish temporal relationships and better understand the mechanisms driving dysbiosis. Additionally, while shotgun metagenomics offers comprehensive insights into microbial functional potential, it does not capture active transcription or metabolic fluxes. Integrating metatranscriptomics or metabolomics in future studies would further elucidate the active processes underlying helminth-induced microbiome disruptions.

In conclusion, this study provides the strongest evidence to date that *T. trichiura* infection leads to both conserved and region-specific disruptions in the gut microbiome, affecting SCFA metabolism, mucin degradation, and microbial network stability. These findings expand previous work by demonstrating that helminth-induced dysbiosis follows a predictable functional trajectory, while also being shaped by local ecological and dietary factors. Understanding these dynamics is critical for developing targeted microbiome-based strategies to mitigate the burden of soil-transmitted helminths and improve gut health in endemic regions.

## Ethical approval and consent to participate

The study protocol was approved by independent ethics committees in Côte d’Ivoire (reference numbers 088–18/MSHP/CNESVS-km and ECCI00918), Laos (reference number 093/NECHR), Pemba (Tanzania, reference number ZAMREC/0003/Feb/2018), and Switzerland (reference number BASEC Req-2018-00494). This trial was registered under the number NCT03527732 on ClinicalTrials.gov. Participants were informed about the purpose of the study and informed consent forms were obtained during information sessions

## Consent for publication

Not applicable

## Availability of data and material

Shotgun sequence data supporting the findings of this study have been deposited in the NCBI Short Read Archive with the primary accession code PRJNAXXXX.

## Author contributions

**PHHS**: study design, research design, project supervision, experimental work, statistical analyses, figure generation, writing of the initial manuscript, manuscript editing; **EH, AS, SS, and JTC**: study design, conducted field work (sample collection, handling, coordination of treatment, parasitological work, and data curation); **JD and NR:** experimental work (DNA isolation, Nanopore sequencing), manuscript editing; **JK**: study design, research design, project supervision, writing of the initial manuscript, manuscript editing.

## Acknowledgements

We are deeply grateful to the dedicated trial team members from the Centre Suisse de Recherches Scientifiques and Université Félix Houphouët-Boigny in Abidjan, Côte d’Ivoire; the Lao Tropical and Public Health Institute in Vientiane, Laos; and the Public Health Laboratory–Ivo de Carneri in Chake Chake, Pemba Island, for their hard work and expertise in both field and laboratory activities. We also extend our heartfelt thanks to the local village and medical authorities in Chake Chake district, Pemba Island; Nambak district, Luang Prabang province, Laos; and the districts of Dabou and Jacqueville in Côte d’Ivoire, whose collaboration and support were invaluable in making this work possible. We thank Christian Beisel from the Genomics Facility Basel for their support during the shotgun sequencing experiments. We are grateful for the access and support to/from the sciCORE (https://scicore.unibas.ch) scientific computing center at the University of Basel.

## Declaration on the use of AI

In the preparation of this manuscript, AI-based tools (ChatGPT and Copilot) were used to assist in language editing, grammatical correction, and improving the clarity of the text. These tools were employed solely for editorial purposes and did not contribute to the conceptualization, data analysis, or scientific conclusions of this work. All scientific content, interpretations, and conclusions presented in this manuscript are the sole responsibility of the authors, who have thoroughly reviewed and approved the final version of the text.

## References

1. Zmora N, Suez J, Elinav E: You are what you eat: diet, health and the gut microbiota. Nature reviews Gastroenterology & hepatology 2019, 16(1):35–56.

2. Levy M, Kolodziejczyk AA, Thaiss CA, Elinav E: Dysbiosis and the immune system. Nature Reviews Immunology 2017, 17(4):219–232.

3. Schneeberger PH, Coulibaly JT, Panic G, Daubenberger C, Gueuning M, Frey JE, Keiser J: Investigations on the interplays between *Schistosoma mansoni*, praziquantel and the gut microbiome. Parasites & vectors 2018, 11:1–12.

4. Leung JM, Graham AL, Knowles SC: Parasite-microbiota interactions with the vertebrate gut: synthesis through an ecological lens. Frontiers in microbiology 2018, 9:843.

5. Hodžić A, Dheilly NM, Cabezas-Cruz A, Berry D: The helminth holobiont: a multidimensional host–parasite–microbiota interaction. Trends in Parasitology 2023, 39(2):91–100.

6. Loke P, Lim Y: Helminths and the microbiota: parts of the hygiene hypothesis. Parasite immunology 2015, 37(6):314–323.

7. Else KJ, Keiser J, Holland CV, Grencis RK, Sattelle DB, Fujiwara RT, Bueno LL, Asaolu SO, Sowemimo OA, Cooper PJ: Whipworm and roundworm infections. Nature Reviews Disease Primers 2020, 6(1):44.

8. Jourdan PM, Lamberton PH, Fenwick A, Addiss DG: Soil-transmitted helminth infections. The lancet 2018, 391(10117):252–265.

9. Lawson MA, Roberts IS, Grencis RK: The interplay between *Trichuris* and the microbiota. Parasitology 2021, 148(14):1806–1813.

10. Reynolds LA, Finlay BB, Maizels RM: Cohabitation in the intestine: interactions among helminth parasites, bacterial microbiota, and host immunity. The Journal of Immunology 2015, 195(9):4059–4066.

11. Schachter J, Alvarinho de Oliveira D, Da Silva CM, de Barros Alencar ACM, Duarte M, da Silva MMP, Ignácio ACdPR, Lopes-Torres EJ: Whipworm infection promotes bacterial invasion, intestinal microbiota imbalance, and cellular immunomodulation. Infection and Immunity 2020, 88(3):10.1128/iai.00642-00619.

12. Chen H, Mozzicafreddo M, Pierella E, Carletti V, Piersanti A, Ali SM, Ame SM, Wang C, Miceli C: Dissection of the gut microbiota in mothers and children with chronic *Trichuris trichiura* infection in Pemba Island, Tanzania. Parasites & Vectors 2021, 14:1–13.

13. Sargsian S, Chen Z, Lee SC, Robertson A, Thur RS, Sproch J, Devlin JC, Tee MZ, Er YX, Copin R: Clostridia isolated from helminth-colonized humans promote the life cycle of *Trichuris* species. Cell reports 2022, 41(9).

14. Tee MZ, Er YX, Easton AV, Yap NJ, Lee IL, Devlin J, Chen Z, Ng KS, Subramanian P, Angelova A: Gut microbiome of helminth-infected indigenous Malaysians is context dependent. Microbiome 2022, 10(1):214.

15. Holm JB, Sorobetea D, Kiilerich P, Ramayo-Caldas Y, Estellé J, Ma T, Madsen L, Kristiansen K, Svensson-Frej M: Chronic *Trichuris muris* infection decreases diversity of the intestinal microbiota and concomitantly increases the abundance of lactobacilli. PloS one 2015, 10(5):e0125495.

16. Hürlimann E, Keller L, Patel C, Welsche S, Hattendorf J, Ali SM, Ame SM, Sayasone S, Coulibaly JT, Keiser J: Efficacy and safety of co-administered ivermectin and albendazole in school-aged children and adults infected with *Trichuris trichiura* in Côte d’Ivoire, Laos, and Pemba Island, Tanzania: a double-blind, parallel-group, phase 3, randomised controlled trial. The Lancet infectious diseases 2022, 22(1):123–135.

17. Bärenbold O, Raso G, Coulibaly JT, N’Goran EK, Utzinger J, Vounatsou P: Estimating sensitivity of the Kato-Katz technique for the diagnosis of *Schistosoma mansoni* and hookworm in relation to infection intensity. PLoS neglected tropical diseases 2017, 11(10):e0005953.

18. Hillmann B, Al-Ghalith GA, Shields-Cutler RR, Zhu Q, Gohl DM, Beckman KB, Knight R, Knights D: Evaluating the information content of shallow shotgun metagenomics. Msystems 2018, 3(6):10.1128/msystems.00069-00018.

19. Blanco-Míguez A, Beghini F, Cumbo F, McIver LJ, Thompson KN, Zolfo M, Manghi P, Dubois L, Huang KD, Thomas AM: Extending and improving metagenomic taxonomic profiling with uncharacterized species using MetaPhlAn 4. Nature Biotechnology 2023, 41(11):1633–1644.

20. Langmead B, Salzberg SL: Fast gapped-read alignment with Bowtie 2. Nature Methods 2012, 9(4):357–359.

21. Lu Y, Zhou G, Ewald J, Pang Z, Shiri T, Xia J: MicrobiomeAnalyst 2.0: comprehensive statistical, functional and integrative analysis of microbiome data. Nucleic Acids Research 2023, 51(W1):W310–W318.

22. Caspi R, Billington R, Keseler IM, Kothari A, Krummenacker M, Midford PE, Ong WK, Paley S, Subhraveti P, Karp PD: The MetaCyc database of metabolic pathways and enzymes-a 2019 update. Nucleic acids research 2020, 48(D1):D445–D453.

23. Team RC: A language and environment for statistical computing. R Foundation for Statistical Computing, *Vienna, Austria* 2024.

24. Oksanen J, Blanchet FG, Kindt R, Legendre P, Minchin PR, O’hara R, Simpson GL, Solymos P, Stevens MHH, Wagner H: Package ‘vegan’. Community ecology package, version 2013, 2(9):1–295.

25. Lahti L, Shetty S, Blake T, Salojarvi J: Tools for microbiome analysis in R. *Version* 2017, 1(5):28.

26. Gu Z: Complex heatmap visualization. Imeta 2022, 1(3):e43.

27. Kurtz ZD, Müller CL, Miraldi ER, Littman DR, Blaser MJ, Bonneau RA: Sparse and compositionally robust inference of microbial ecological networks. PLoS computational biology 2015, 11(5):e1004226.

28. Gustavsen JA, Pai S, Isserlin R, Demchak B, Pico AR: RCy3: Network biology using Cytoscape from within R. F1000Research 2019, 8.

29. Csardi G, Nepusz T, Traag V, Horvát S, Zanini F, Noom D, Müller K: Igraph: Network analysis and visualization. R package version 2023, 1(0.9002).

30. Rupert Jr G: Simultaneous statistical inference. 2012.

31. Benjamini Y, Hochberg Y: Controlling the false discovery rate: a practical and powerful approach to multiple testing. Journal of the Royal statistical society: series B (Methodological) 1995, 57(1):289–300.

32. Segata N, Izard J, Waldron L, Gevers D, Miropolsky L, Garrett WS, Huttenhower C: Metagenomic biomarker discovery and explanation. Genome biology 2011, 12:1–18.

33. Mallick H, Rahnavard A, McIver LJ, Ma S, Zhang Y, Nguyen LH, Tickle TL, Weingart G, Ren B, Schwager EH: Multivariable association discovery in population-scale meta-omics studies. PLoS computational biology 2021, 17(11):e1009442.

34. Organization WH: Assessing schistosomiasis and soil-transmitted helminthiases control programmes: monitoring and evaluation framework. 2024.

35. Koh A, De Vadder F, Kovatcheva-Datchary P, Bäckhed F: From dietary fiber to host physiology: short-chain fatty acids as key bacterial metabolites. Cell 2016, 165(6):1332–1345.

36. Bethony J, Brooker S, Albonico M, Geiger SM, Loukas A, Diemert D, Hotez PJ: Soil-transmitted helminth infections: ascariasis, trichuriasis, and hookworm. The lancet 2006, 367(9521):1521-1532.

37. Maizels RM, Smits HH, McSorley HJ: Modulation of host immunity by helminths: the expanding repertoire of parasite effector molecules. Immunity 2018, 49(5):801–818.

38. Donaldson GP, Lee SM, Mazmanian SK: Gut biogeography of the bacterial microbiota. Nature Reviews Microbiology 2016, 14(1):20–32.

39. Ramanan D, Bowcutt R, Lee SC, Tang MS, Kurtz ZD, Ding Y, Honda K, Gause WC, Blaser MJ, Bonneau RA: Helminth infection promotes colonization resistance via type 2 immunity. Science 2016, 352(6285):608-612.

