## Supplementary figures for "Profound taxonomic and functional gut microbiota alterations associated with trichuriasis: cross-country and country-specific patterns"

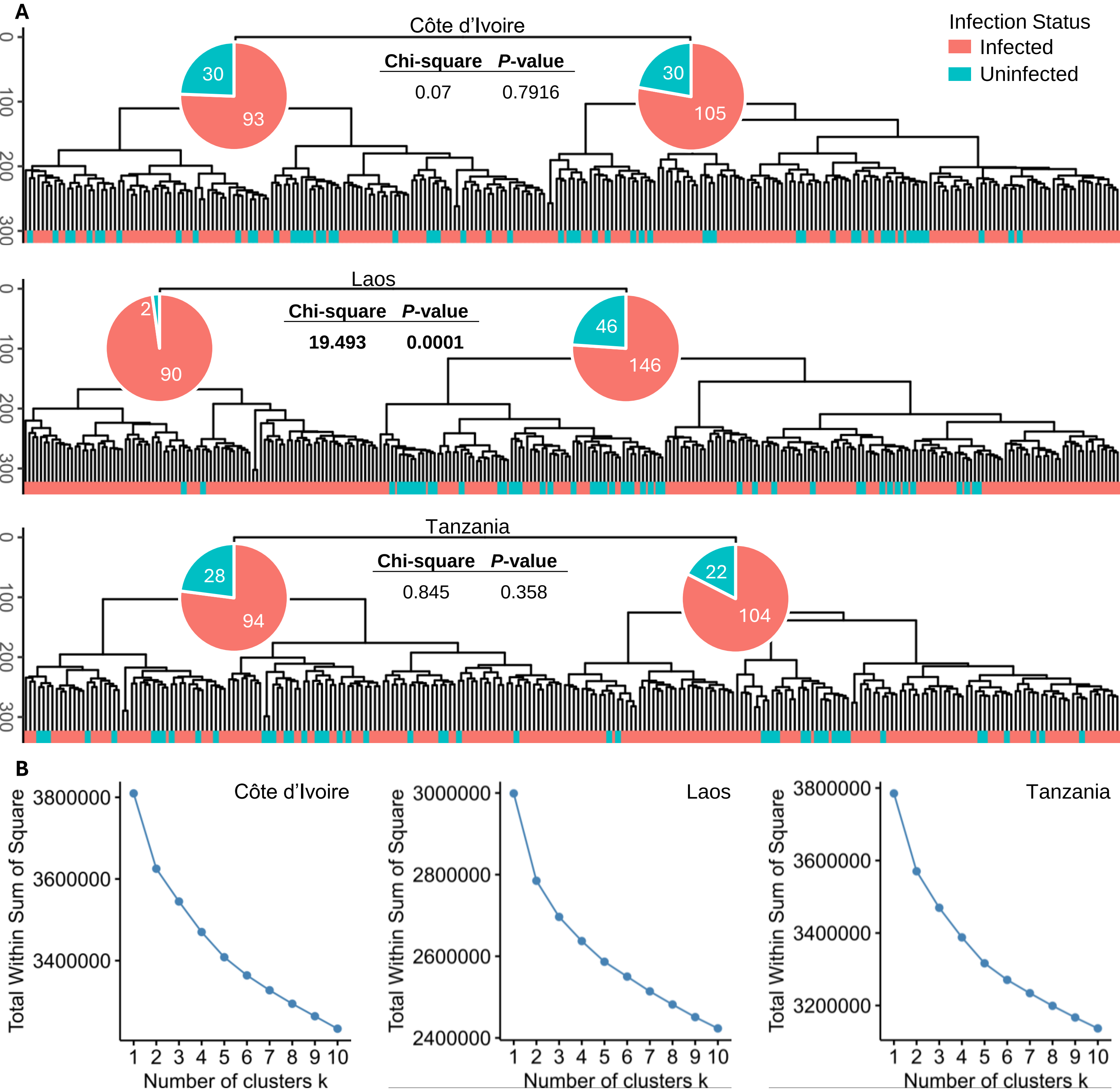


**Supplementary Figure 1. Panel A.** Clustering of samples based on their species composition, by country. Clustering was done on centered log ratio transformed species-level taxonomic abundance tables using Euclidean distance and the Ward.D2 algorithm. **Panel B.** Elbow method was performed using the within sum of squares approach to analyze within cluster variance and determine the optimal number of clusters in each country.

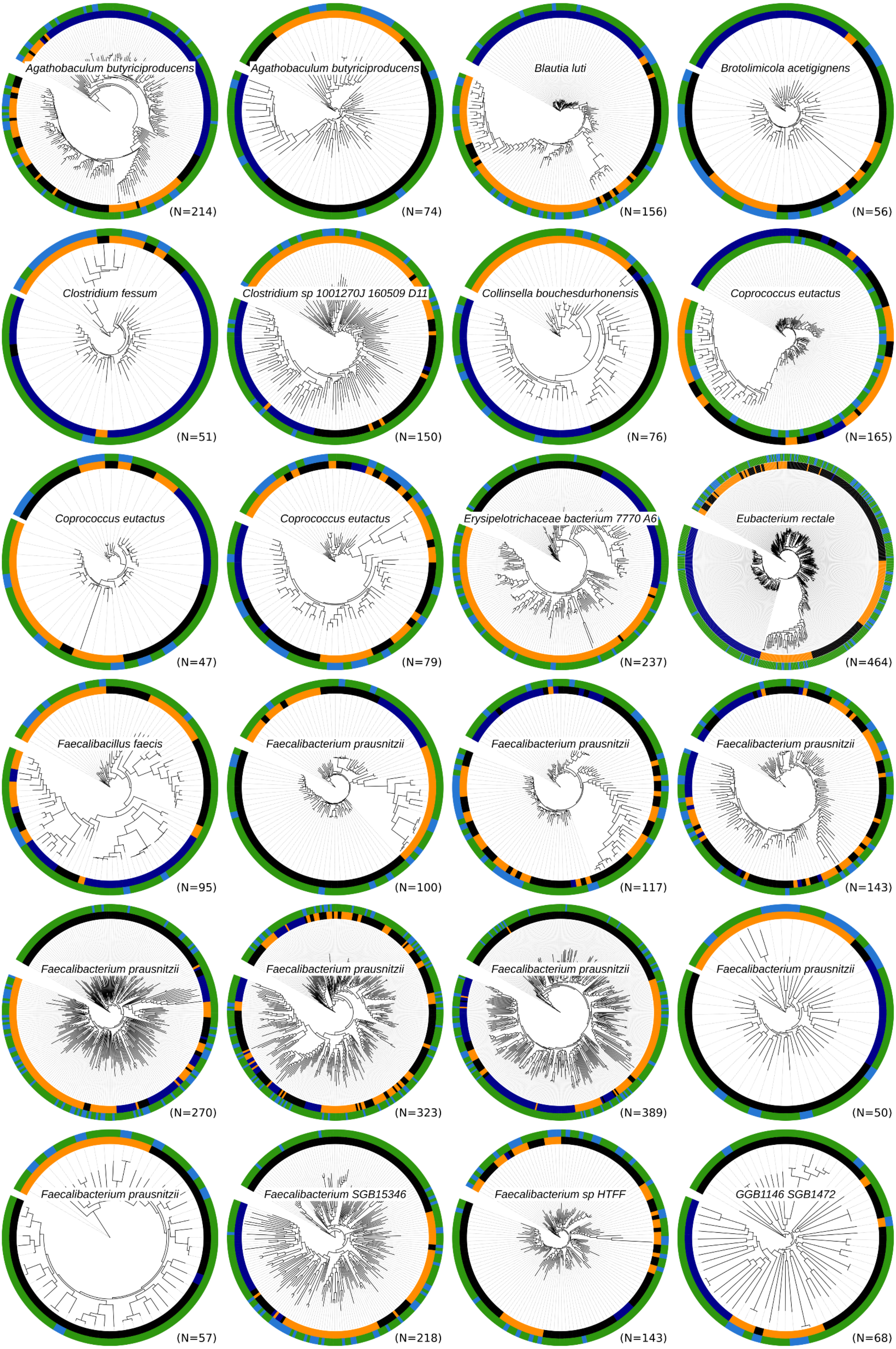
Agathobaculum butyriciproducens
(N=214)
Agathobaculum butyriciproducens
(N=74)
Blautia luti
(N=156)
Brotolimicola acetigignens
(N=56)
Clostridium fessum
(N=51)
Clostridium sp 1001270J 160509 D11
(N=150)
Collinsella bouchesdurhonensis
(N=76)
Coprococcus eutactus
(N=165)
Coprococcus eutactus
(N=47)
Coprococcus eutactus
(N=79)
Erysipelotrichaceae bacterium 7770 A6
(N=237)
Eubacterium rectale
(N=464)
Faecalibacillus faecis
(N=95)
Faecalibacterium prausnitzii
(N=100)
Faecalibacterium prausnitzii
(N=117)
Faecalibacterium prausnitzii
(N=143)
Faecalibacterium prausnitzii
(N=270)
Faecalibacterium prausnitzii
(N=323)
Faecalibacterium prausnitzii
(N=389)
Faecalibacterium prausnitzii
(N=50)
Faecalibacterium prausnitzii
(N=57)
Faecalibacterium SGB15346
(N=218)
Faecalibacterium sp HTFF
(N=143)
GGB1146 SGB1472
(N=68)

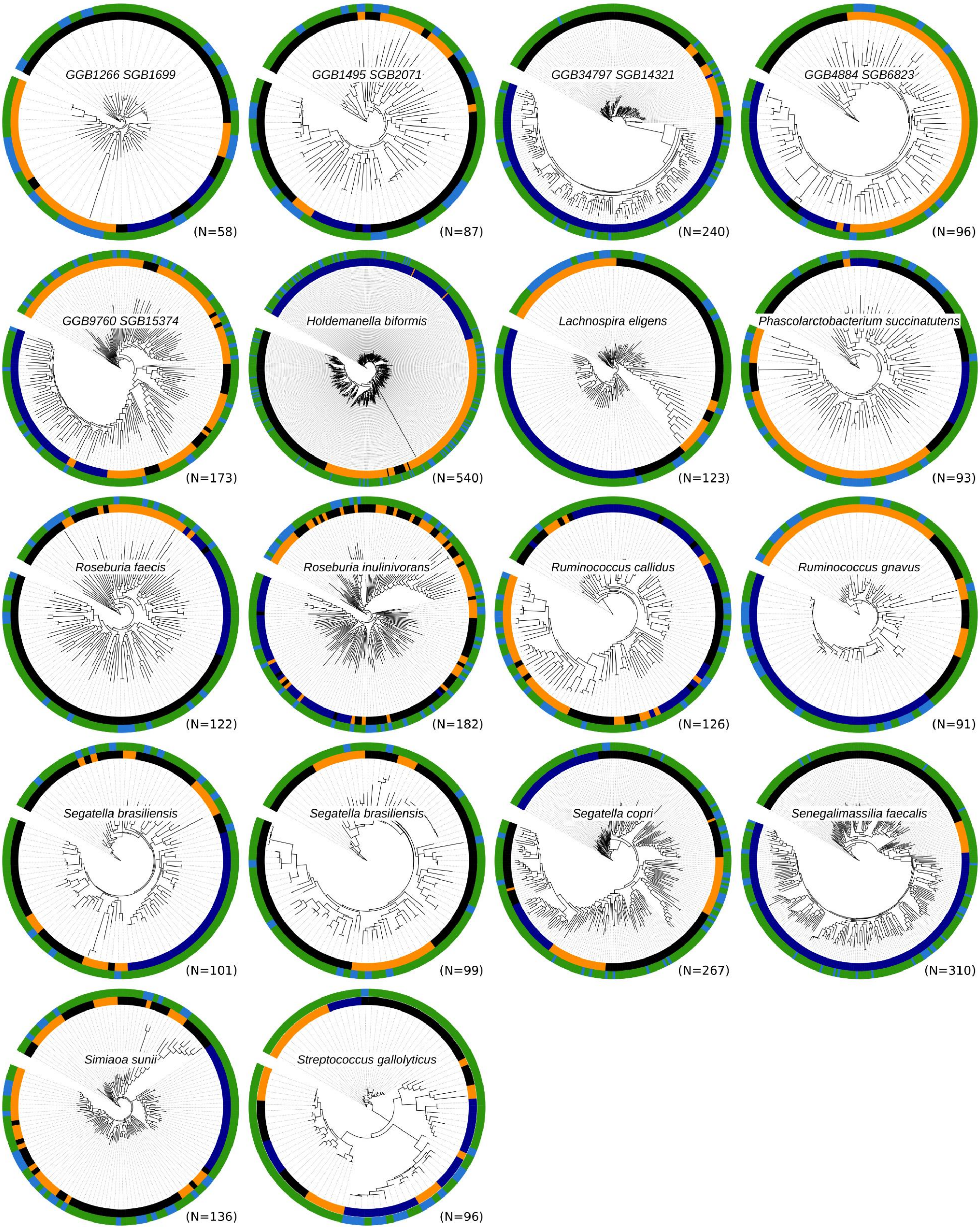


**Supplementary Figure 1. Panel A.** Clustering of samples based on their species composition, by country. Clustering was done **Panel B.** Elbow method to analyze the within cluster variance to determine the optimal number of clusters.

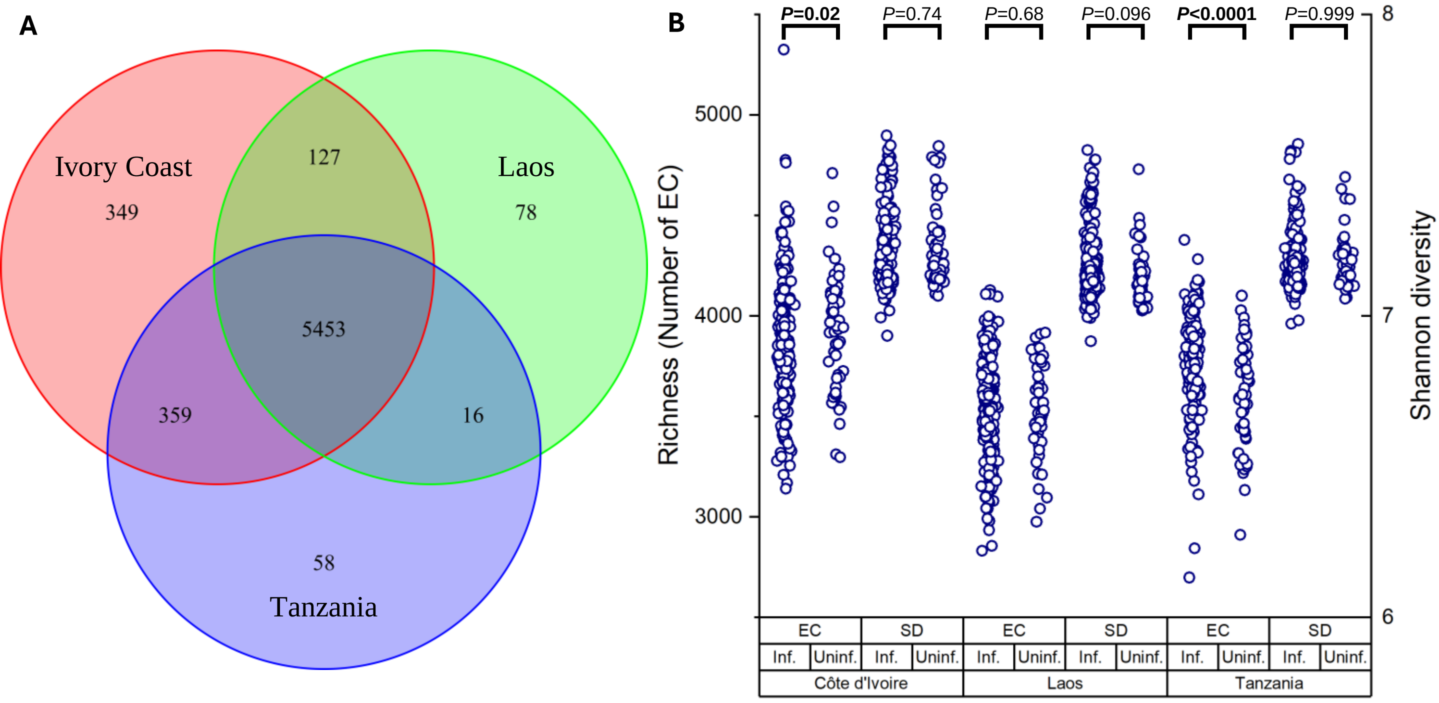


**Supplementary Figure 3. Panel A.** Venn diagram showing the number of common and unique enzyme commission numbers (EC) found in all samples. **Panel B.** Comparison of the functional diversity, including richness (Number of ECs) and Shannon diversity index, stratified by country and infection status.
